# A self-assembling cross-protective antigen against multiple Gram-positive nosocomial pathogens

**DOI:** 10.1101/2025.02.06.636879

**Authors:** Eliza Kramarska, Felipe Romero-Saavedra, Flavia Squeglia, Sara La Manna, Oceane Sadones, Daniela Marasco, Rita Berisio, Johannes Huebner

## Abstract

ESKAPE pathogens are responsible for complicated nosocomial infections worldwide and are usually resistant to commonly used antibiotics in clinical settings. Among these bacteria, vancomycin-resistant *Enterococcus faecium* and methicillin-resistant *Staphylococcus aureus* are the two most important Gram-positive pathogens for which alternative treatments and preventions are urgently needed. We previously designed a multi-presenting antigen, embedding the main epitope displayed by the AdcA protein of *E. faecium*, that conferred protection against different Gram-positive pathogens both in passive and active immunization models. Here, we developed a new presentation strategy for this epitope, the EH-motif, based on a self-assembling peptide. Self-assembling peptides have been promising in the fields of material sciences, nanoscience, and medicine and have also potential in vaccine development, as they allow multiple presentations of the epitope and provide an ideal size for production and application. We show that this multi- presenting peptide, here Q11-EH, forms stable fibers of nanometric size. We also demonstrate that antibodies raised against Q11-EH mediate the opsonic killing of a wide-spectrum of Gram-positive pathogens, including *E. faecium*, *S. aureus,* and *E. faecalis*. Our data indicate that multiple presentation strategies are a potent tool for vaccine antigen improvement and point to Q11-EH as a promising antigen for the development of novel cross-protective vaccines.

## Introduction

Antimicrobial resistance (AMR) represents an environmental adaptation, which allows pathogens to remain active in the presence of bacteriostatic or bactericidal concentrations of antimicrobials (1). Recently, bacterial resistance has been identified by the WHO as one of the 10 biggest threats in healthcare for the next decades. (2). ESKAPE bacteria, including *Enterococcus faecium, Staphylococcus aureus, Klebsiella pneumoniae, Acinetobacter baumanii, Pseudomonas aeruginosa, Enterobacter* spp, and *Escherichia coli*, are common etiological agents of Nosocomial Infections (Nis) (3), and leading causes of AMR-associated deaths (4). These bacteria are usually resistant to the majority of standard treatments, are able to quickly adapt to newly developed strategies, and exhibit resistance to important antimicrobials (5). Particularly dangerous are phenotypes of gram-negative bacteria producing extended-spectrum beta-lactamases (ESBLs), and of Gram-positive bacteria with resistance to broad-spectrum last-resort antibiotics, such as vancomycin-resistant *E. faecium* (VRE) (6) and methicillin-resistant *S. aureus* (MRSA).

Enterococci are ubiquitous Gram-positive, facultative anaerobic microorganisms, which colonize a broad range of hosts, from invertebrates to mammals, including humans (7, 8). They are among the most important pathogens of NIs and within this genus, *E. faecium* and *E. faecalis* have the highest clinical relevance (9, 10) infecting patients with recent surgery, organ transplantation, diabetes, malignancy, and renal insufficiency (11, 12). Although both species have clinical importance, *E. faecium* infections have higher rates of antibiotic resistance and mortality (13). Already in 2017, vancomycin-resistant *E. faecium* has been declared by WHO a threat to humankind, for which rapid actions are needed (14),(15). With the rise of vancomycin resistance, new treatments were developed and introduced but this accelerated the selection of multidrug or even pan-resistant enterococci (13, 16–20). Similarly, *S. aureus* can be easily found in the environment, on the skin, and on mucous membranes, especially in the nares, where it is present among 25-30% of healthy individuals. However, despite being a commensal (21), *S. aureus* is also one of the most common causative agents of bacterial toxin-mediated diseases and invasive infections, as it can lead to bacteremia, endocarditis, skin and soft tissue infections, septic arthritis, prosthetic device infections, pulmonary tract infections, gastroenteritis, meningitis, toxic shock syndrome and urinary tract infections (22). MRSA and VRE represent a serious concern within hospitals and long-term care units, posing a threat to patients of all ages.

Despite several proposed candidate vaccine antigens against *E. faecium* (23–27), the WHO report from 2021 indicated a lack of vaccines against *E. faecium* in preclinical development (28). Similarly, no vaccine against *S. aureus* is currently available (28), although diverse and complex candidates targeting this pathogen are under investigation (29, 30). In a previous work, we used a structural vaccinology approach to develop a vaccine antigen preparation that is cross-reactive against several Gram-positive pathogens (31). To achieve this goal, we identified a promising antigenic conserved region of the key zinc transporting lipoprotein AdcA of *E. faecium,* which we denominated “EH-motif” (31). AdcA, a 508 residues and multi-domain protein, was shown to elicit specific, opsonic, protective antibodies, with extensive coverage among the homologous strain *E. faecium* E155 and several clinical *E. faecium* isolates (23, 32). Based on this finding, we designed and developed a multi-presenting antigen that carried three EH-motifs on each molecule. This antigen, Sc(EH)_3_, proved to elicit opsonic and protective antibodies that are effective against Gram-positive pathogens, including *E. faecium, E. faecalis and S. aureus* (31). The immune-stimulating properties of the EH-motif and the multiple presentations in Sc(EH)_3_ prompted us to design diverse effective multiple-presentation strategies for the EH-motif (31).

Self-assembling peptides provide several advantages in biomedical applications, including multi- valency and ease of synthetic modification (33–36). Among them, the short fibrillising peptide, Q11 (AcQQKFQFQFEQQ-Am) proved to self-assemble in salt-containing aqueous environments to form networks of β-sheet-rich nanofibers (34). Also, the addition of cell-binding amino acid sequences to the N terminus of Q11 led to self-assembled fibrils that functionally present the cell-binding peptides on their surfaces (37). Furthermore, Q11 and other self-assembling peptide-based materials have been found to be only minimally immunogenic (37, 38).

In the present work, we sought to design a supra-molecular vaccine antigen, where our previously identified strong AdcA-derived epitope, the EH-motif (31), is displayed in multiple copies on the Q11-based nanofiber (37). We synthesized a modified Q11 peptide, by adding the EH-motif at its N-terminus, separated by a short peptide linker. Polymerization of this molecule, here designated Q11-EH, led to nanostructures of 229 nm average size. Our results indicate a surprisingly strong antibody response generated against Q11-EH. Most importantly, Q11-EH elicits opsonic antibodies in rabbits that are effective against multiple Gram-positive pathogens, including *E. faecium, E. faecalis*, and *S. aureus*.

## RESULTS

### Design and characterisation of the epitope multi-presenting antigen Q11-EH

A self-assembling molecule carrying our previously identified EH-motif of the AdcA lipoprotein from *E. faecium* was designed to exploit multiple antigen presentations. To achieve this, we started from the previously reported Q11 peptide and added the sequence of the EH motif at its N-terminal end, separated by a short peptide linker (SGSG) as a flexible spacer between the Q11 scaffold and the antigenic EH-motif (Figure 1). Q11-EH was synthesized through manual solid-phase peptide synthesis to high purity, as evaluated by LC-MS analysis (Figure S1).

**Figure 1.**
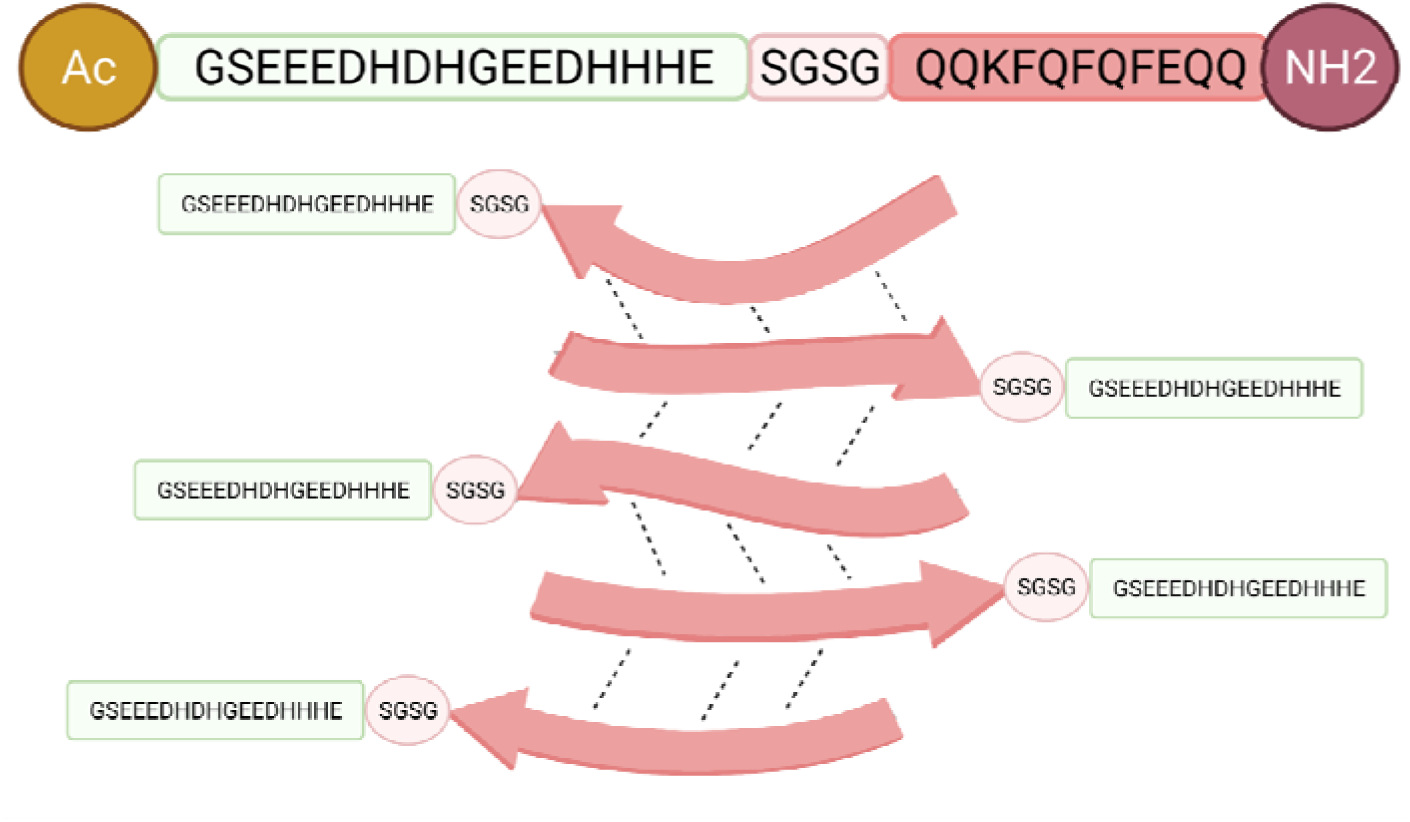
Graphical representation of the Q11-EH multi-presenting antigen. The EH-motif is represented in green, the short linker in light pink, and the Q11 peptide in dark pink. The peptide is acetylated at its N-terminal end and amidated at its C-terminal end.

As previously reported, salts are known to start the polymerization of Q11 in a time-dependent manner (34). Therefore, we resuspended Q11-EH in water to obtain a 10mM stock solution and then prepared several dilutions with peptide concentrations ranging from 1 µM to 125 µM, in PBS at pH 7.4 to establish a polymerization protocol, through the time-dependent examination of the supramolecular dimensions using Dynamic Light Scattering (DLS). We observed that polymerization was a concentration-dependent process in the first days. However, the degree of monodispersion increased with time and all samples reached the same dimensions after 5 days of storage at 4°C. Using Q11-EH 1 µM concentration, polymerization was complete after 96 hours, when the sample was 100% monodisperse (Figure 2 and Table 1), with a hydrodynamic (Z-average) radius of (2.3±0.3)*10^2^ nm. No changes in the size distribution were measured by DLS for longer time storage (Table 1). The estimated molecular weight for the stable polymer, (1.1 ± 0.2)*10^6^, indicates that a single fibril contains approximately 300 copies of the Q11-EH peptide.

**Figure 2.**
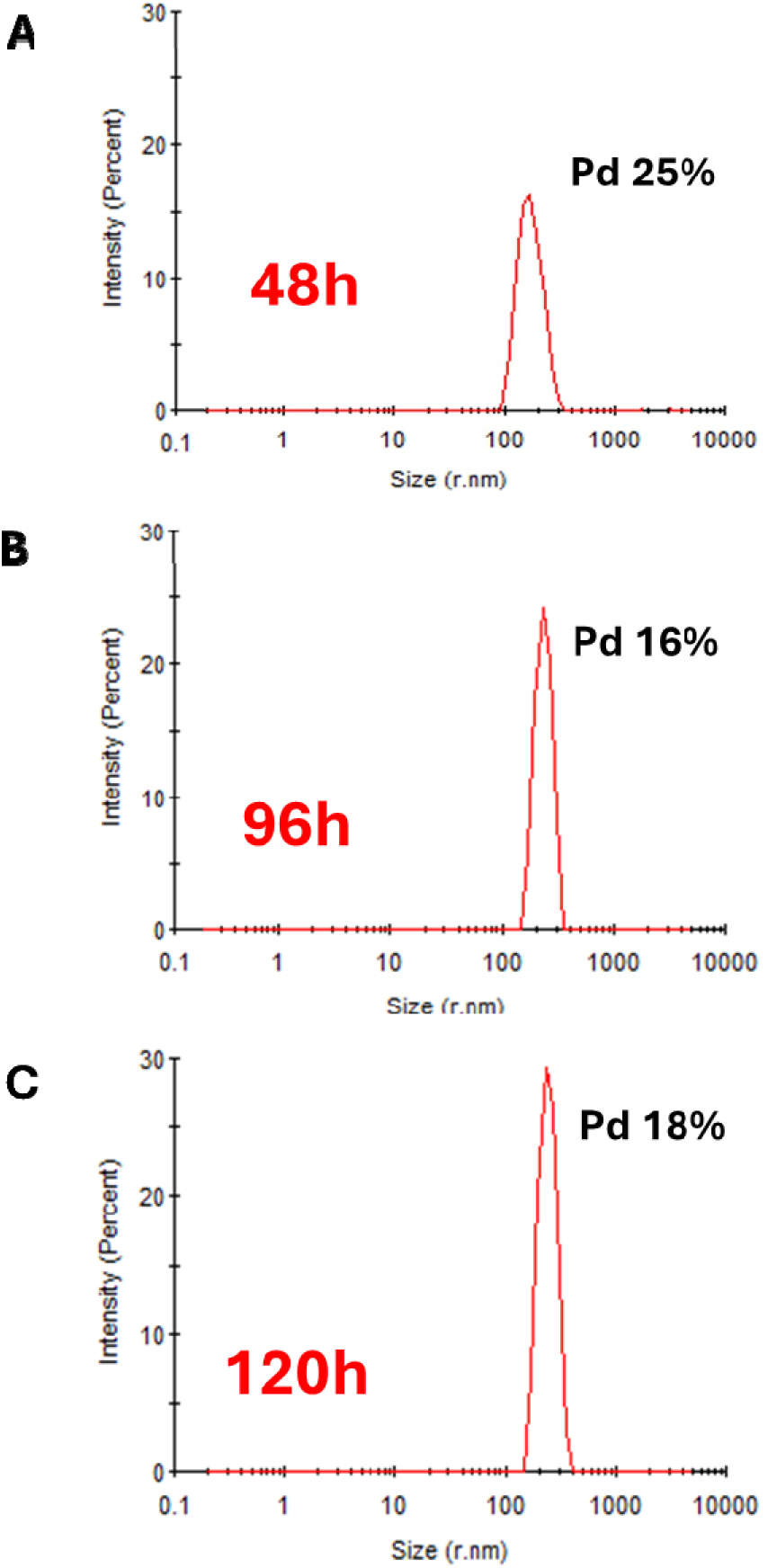
(A-C) Size distribution of Q11-EH particles by DLS during the polymerization process. Pd is the polydispersity index. At 96h the sample is fully monodisperse and the hydrodynamic radius remains constant with time. A- 48h, B – 96h, C – 120h of polymerization.

**Table 1.** Dependence of the hydrodynamic radius on the polymerization time. The Z-average radius corresponding to each peak was calculated from the correlation function using the Malvern technology software.

| Time (h) | Hydrodynamic radius (nm) | Estimated Mw (kDa) | Pd (%) | Polydispersity |
| --- | --- | --- | --- | --- |
| 48 | $(1.7 \pm 0.4) \cdot 10^2$ | $(5.6 \pm 1.4) \cdot 10^5$ | 25 | Polydisperse |
| 96 | $(2.3 \pm 0.3) \cdot 10^2$ | $(1.1 \pm 0.2) \cdot 10^6$ | 16 | Monodisperse |
| 120 | $(2.3 \pm 0.3) \cdot 10^2$ | $(1.1 \pm 0.2) \cdot 10^6$ | 18 | Monodisperse |

As further proof that the formed polymeric species adopt a β-sheet-rich structure like its progenitor peptide Q11 (34), we used the Thioflavin T (ThT) assay (39). As shown in Figure 3, we observed an increase in fluorescence emission over time when in complex with Q11-EH, a feature which is typical of β-sheet-rich fibrils.

**Figure 3.**
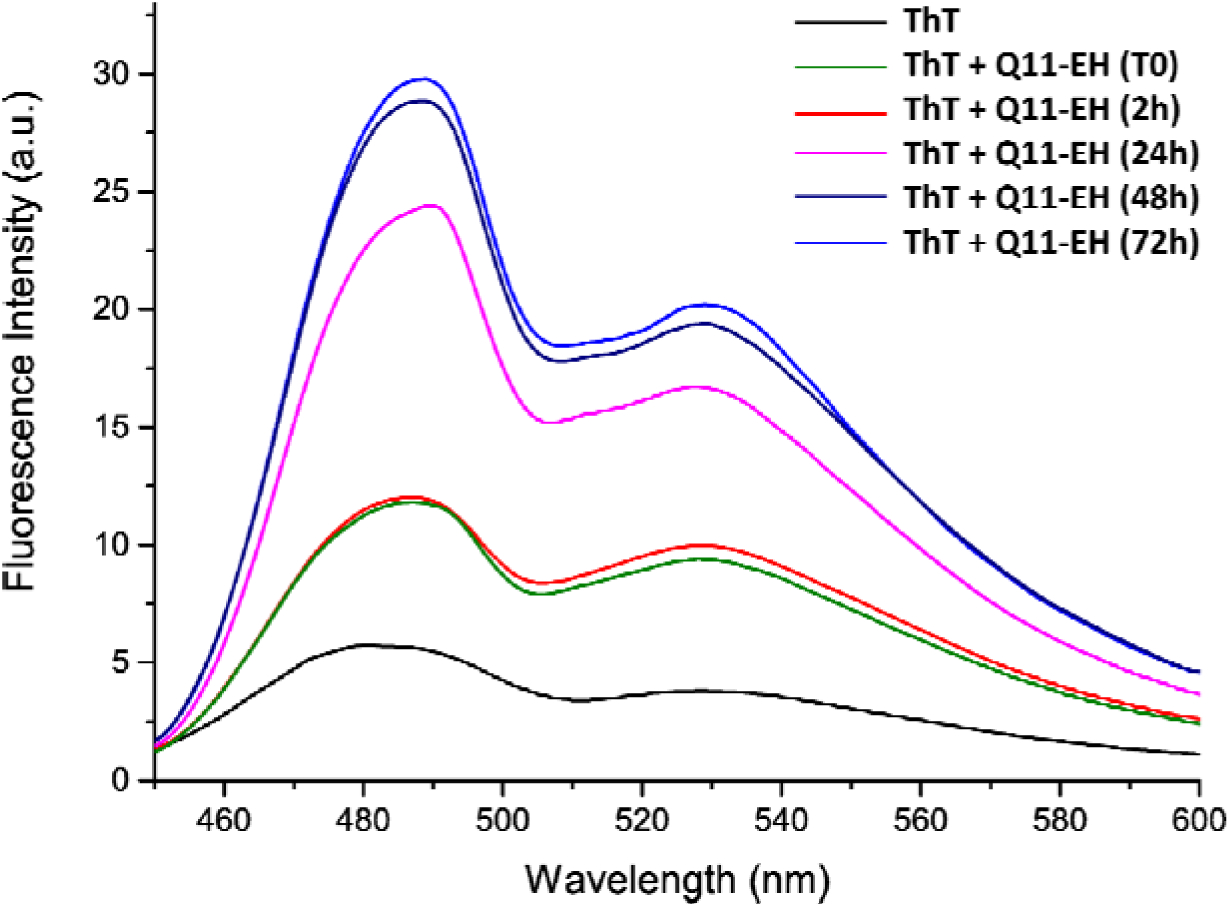
Time course of ThT fluorescence emission intensity when in complex with Q11-EH. Overlaid spectra at the specified time points are presented. Fluorescence was reported as Arbitrary Unit after excitation at 440 nm and recorded between 450 and 600 nm.

### Q11-EH supra-structure elicits antibodies able to induce bacterial killing

After the polymerization protocol, polymerized Q11-EH was used to immunize two New Zealand white rabbits, which were exsanguinated two weeks after the last injection. To generate a serum that represents an average immune response, the terminal bleeds of rabbits vaccinated with the same compound were mixed in equal volumes, generating the pooled anti-Q11-EH sera. The same procedure was used for the pre-immune sera used as controls. The opsonophagocytic killing assay (OPA) is used as a surrogate marker to assess whether antibodies are able to mediate a protective immune response (40) (Figure S2). We used this assay to test if anti-Q11-EH antibodies can mediate the opsonic killing of a panel of a panel of Gram-positive bacteria. As shown in Figure 4, anti-Q11-EH antibodies induced *E. faecium* opsonic killing significantly better than antibodies raised against the full-length AdcA. Indeed, an average 67%, 52% and 42% of killing was induced by anti-Q11-EH antibodies 0.4 mg/mL, 0.2 mg/mL, and 0.1 mg/mL, respectively. Opsonic killing induced by these concentrations of anti-AdcA was 39%, 31% and 23%, respectively (Figure 4).

**Figure 4.**
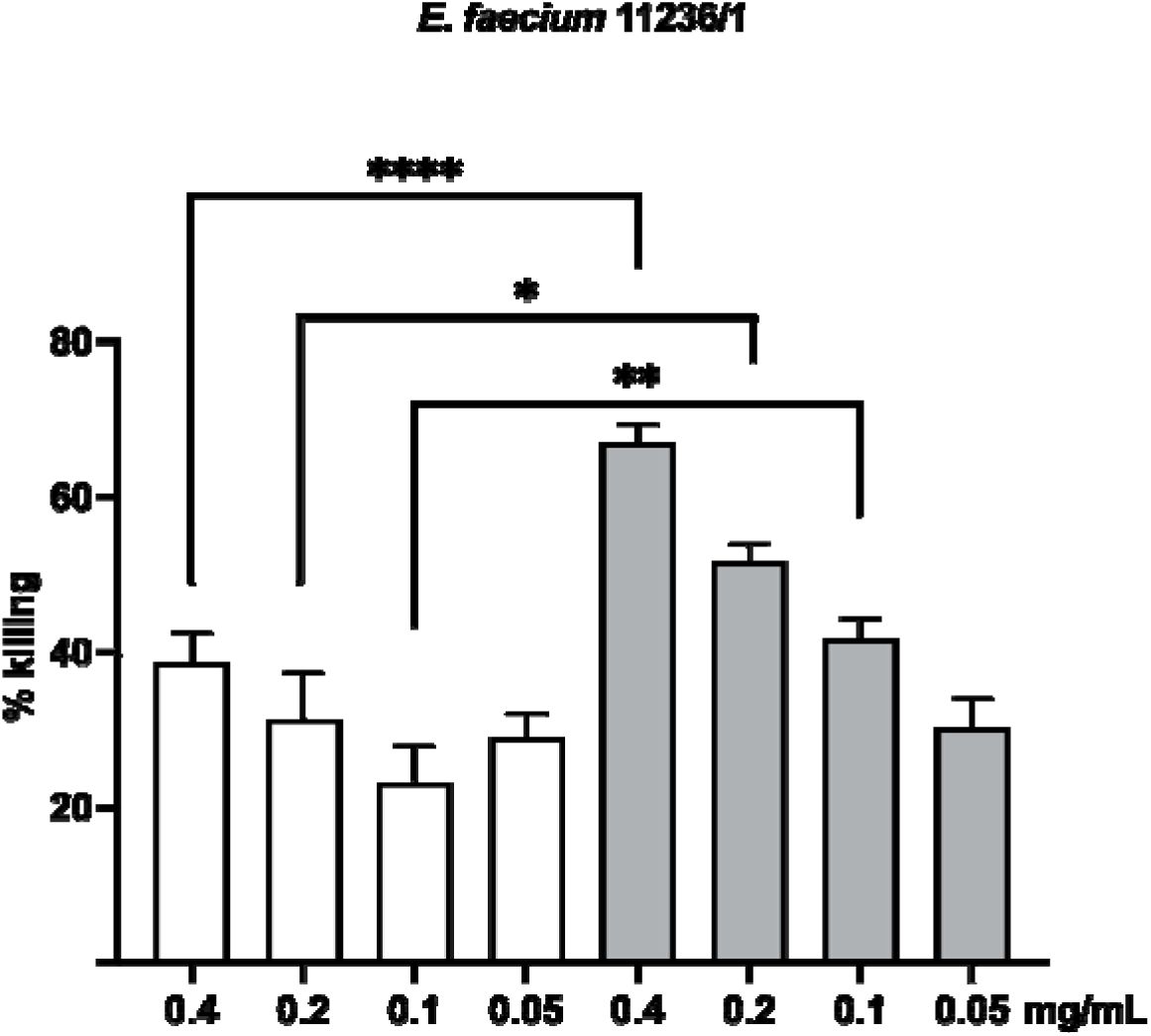
OPA against *E. faecium* 11236/1. Killing percentages mediated by anti-Q11-EH IgG (light grey bars) were compared to those mediated by anti-AdcA (white bars). The statistical significance was tested by the unpaired two-tailed Welch’s t-test with a 95% confidence interval and Bonferroni correction, using anti-AdcA and anti-Q11-EH IgG at the same concentration. Bars and whiskers denote mean values ± standard errors of the mean. *P ≤ 0.0125, ** P ≤0.01, *** P≤0,001.

Since we previously showed that the EH-loop is strongly conserved in Gram-positives (31), we extended this study to *S. aureus* MW2, and *E. faecalis* T2. In the assay for *S. aureus* MW2, all tested concentrations performed significantly better than anti-AdcA sera with bacteria-killing of approximately 66%, 48%, 38%, and 29% for 0.4 mg/mL, 0.2 mg/mL, 0.1 mg/mL and 0.05 mg/mL of IgG (Figure 5A). These values are to be compared to 46%, 32%, 19% and 3% killing induced by anti-AdcA, showing that the best performance of anti-Q11-EH antibodies against *S. aureus* MW2 is observed at the lowest concentration, with 10-fold enhancement of opsonic killing (Figure 5A). Strong, albeit lower, opsonic killing was observed against *E. faecalis* T2 (Figure 5B), reaching 57%, 33%, 15%, 6% (compared to 44%, 22%, 11%, 0% for anti-AdcA), for concentrations 0.4 mg/mL, 0.2 mg/mL, 0.1 mg/mL, and 0.05 mg/mL, respectively.

**Figure 5.**
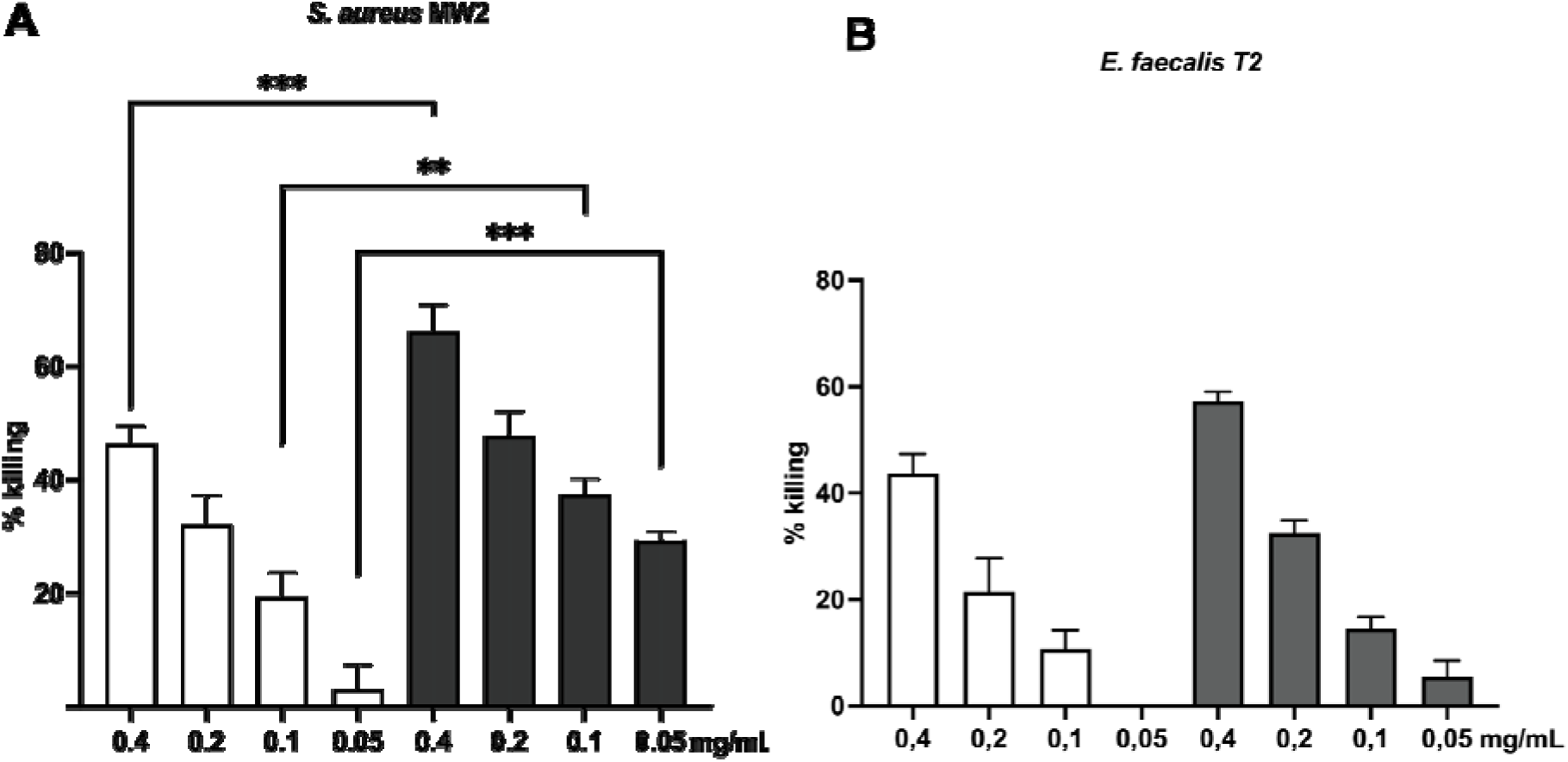
OPA against *S.aureus* MW2 (A) and *E. faecalis* T2(B). Killing percentages mediated by anti-Q11-EH IgG (dark grey bars) were compared to those mediated by anti-AdcA (white bars). Statistical significance was assessed using an unpaired two-tailed Welch’s t-test with a 95% confidence interval and Bonferroni correction. For non-normally distributed data (S. aureus at 0.4 and 0.2 and E. faecalis at 0.2), the Mann-Whitney test with Bonferroni correction was applied. All comparisons were performed using anti- AdcA and anti-Q11-EH IgG at the same concentration. Bars and whiskers represent mean values ± standard error of the mean (SEM). *P ≤ 0.0125, ** P ≤0.01, *** P≤0,001.

### Assessment of antibody specificity using opsonophagocytic inhibition assay

The specificity of opsonising antibodies against the Q11-EH was assessed using opsonophagocytic inhibition assays, by pre-incubating the anti-Q11-EH sera with different inhibitors at different concentrations. Specifically, 0.4 mg/mL anti-Q11-EH sera were incubated with Q11-EH, recombinant AdcA, recombinant ZnuA (the AdcA domain containing the EH epitope) in concentrations ranging between 3.56 and 0.14 µM. In all experiments, we observed a concentration-dependent reduction of killing, suggesting an inhibiting interaction of antibodies with all tested molecules. We observed that 3.56 µM and 1.78 µM of all inhibitors were able to significantly decrease killing. In OPIA with *E. faecium* 11236/1, the highest concentrations (3.56 µM) of Q11- EH, AdcA, ZnuA and reduced killing by 54%, 42% and 42%, respectively (Figure 6A). A similar trend was observed in OPIA with *S. aureus* MW2, where the incubation of anti-Q11-EH with maximum concentrations of Q11-EH, AdcA and ZnuA reduced killing by 65%, 55% and 59%, respectively (Figure 6B). In *E. faecalis* T2, reduction of killing due to the presence of 3.56 µM of Q11-EH, AdcA and ZnuA reached 81%, 77% and 95% respectively (Figure 6C). All these experiments confirmed the specific recognition of anti-Q11-EH antibodies for the EH-motifs of all inhibitors.

**Figure 6.**
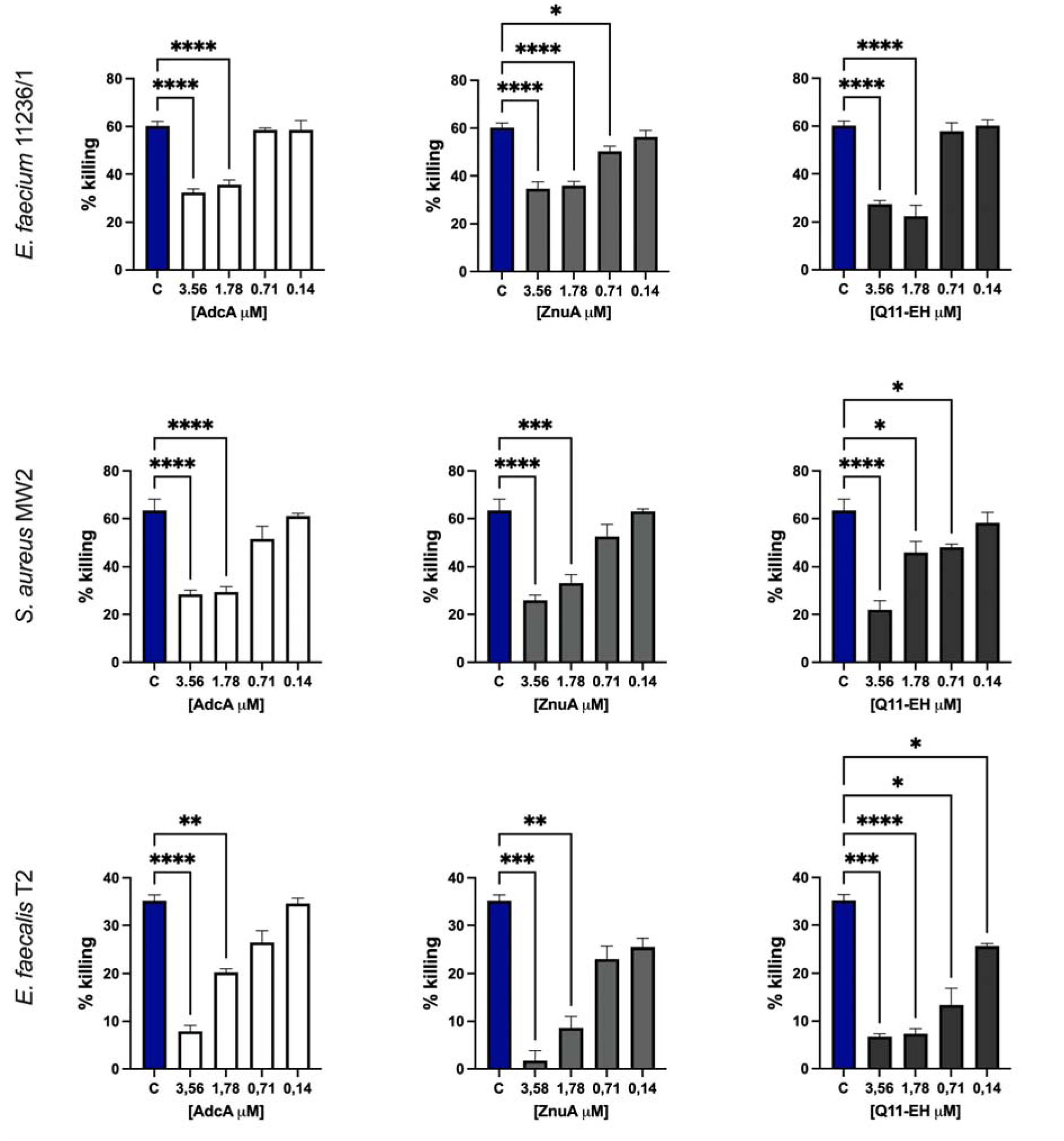
OPIA assay with 0.4 mg/mL of anti-Q11-EH sera against (A) *E. faecium* 11236/1 (B) *S.aureus* MW2 and (C) *E. faecalis* T2. As a killing control “C” for each plot, we used sera at 0.4 mg/mL anti-Q11-EH (blue) without inhibitors. Anti-Q11-EH were incubated with different concentrations of inhibitors, including AdcA (white), ZnuA (light grey), and Q11-EH (dark grey). The final inhibitor concentration is shown below the x-axis expressed as µM. Statistical significance of inhibition was performed by the One-way analysis of variances test, followed by Dunnett’s multiple comparison post hoc test. Bars and whiskers denote mean values ± standard errors of the mean. *P ≤ 0.05, ** P ≤0.01, *** P≤0,001.

## Material and methods

### Design and synthesis of Q11-EH

EH-antigen from AdcA was added to the sequence of a fibrillating peptide, with sequence QQKFQFQFEQQ (Q11) with an addition of a small linker to provide better exposure for the epitope. To the terminal ends acetylation and amidation of the N- and C-terminal ends were added, respectively. Q11-EH peptide was synthesized through manual solid-phase peptide synthesis with a 20 μM scale, following the Fmoc strategy and using standard Fmoc-derivatized amino acids and RINK AMIDE resin with substitution 0.3 mmol/g, as solid support. Activation of amino acids was carried out using HATU and DIEA, whereas Fmoc deprotection with a 20% (v/v) piperidine solution in DMF. All couplings were performed for 20 minutes and deprotections for 10 minutes. After assembly, a small amount of crude product was detached from the resin by treatment with a TFA:TIS:H_2_O (90:5:5 v/v/v) mixture for 1.5 h at room temperature, then it was precipitated in cold ether, dissolved in an H_2_O/CH_3_CN mixture and analyzed by LC-MS analysis (Figure 1). The main peak corresponded to 3780.40 a.m.u. is in perfect agreement with the theoretical one of 3779.69 a. m. u. Then the peptide was acetylated, detached from the resin (TFA:TIS:H_2_O (90:5:5 v/v/v), for 3h), and lyophilized. It was purified by preparative RP-HPLC, applying a linear gradient of 0.1% TFA CH_3_CN in 0.1% TFA water from 5-70% over 13 min with a semipreparative 2.2 × 5 cm C18 column at a flow rate of 20 mL/min, using a UV detector set at a wavelength of 210 nm. The purity of collected fractions was evaluated by LC-MS analysis (Figure S1).

### Dynamic Light Scattering

Polymerization of Q11-EH was established by performing Dynamic Light Scattering (DLS) experiments using a Malvern NanoZetasizer (Malvern, UK). Q11-EH was diluted in Phosphate-Buffered Saline (PBS) at pH 7.4 to a concentration of 1 µM, 32 µM, and 125 µM from a 10 mM stock solution in MilliQ water. Afterward, the samples were sonicated and then incubated at 4°C with agitation. The rate of aggregation was monitored from time 0 to 6 days of incubation. For the analysis of the oligomeric state, the samples were separately loaded into a disposable cuvette and maintained at 20°C during analysis. Spectra were recorded three times with 12 sub-runs using the multimodal mode. Samples with polydispersity index (Pd) lower than 25% are considered monodisperse. The hydrodynamic diameter corresponding to the monodisperse peak was calculated from the correlation function using the Malvern technology software.

### Thioflavin T assay

Thioflavin T (ThT) assay was used to confirm the presence of β-sheets formation in Q11-EH supramolecules. ThT from a stock solution 5mM was diluted to a final concentration of 100µM and mixed with 500µM of Q11-EH in PBS pH 7.4. Fuorescence spectra over the time were collected at 20°C using a Spectrofluorometer Jasco FP-8350 and a quartz cell of 10 mm path-length. ThT fuorescence emission spectra were acquired in the range 450 600 nm upon excitation at 440 nm.

### Production of rabbit polyclonal sera

The immunization protocol using the Q11-EH antigen was conducted by Biogenes GmbH (Berlin, Germany), as outlined in the reference (31). The procedure adhered to national and international animal welfare guidelines for the housing, immunization, and serum collection of rabbits. The protocols were approved by the National Institutes of Health Office of Laboratory Animal Welfare (identifier A5755-0)

### IgG quantification

The concentration of IgG antibodies was determined using the sandwich ELISA method(41). Plates were coated overnight with anti-rabbit IgG at 1 µg/mL concentration in 0,2 M sodium carbonate/bicarbonate buffer, pH=9.4, and kept at 4°C. Blocking was carried out using 3% BSA in PBS solution for 1h at room temperature (RT). Investigated sera and rabbit IgG used for calibration curve were diluted in 1% BSA, 0,5% Tween20, PBS solution and incubated on the plate for 2h at RT. For detection of anti-rabbit, IgG Ab conjugated AP, was likewise incubated for 2h at RT. All the steps were separated by washing in 0.9% NaCl solution with 0.1% Tween20. For the detection disodium p-nitrophenyl phosphate was used at a final concentration of 1 mg/mL in 0.1 M glycine buffer with 1mM MgCl_2_ and 1mM ZnCl_2_, pH 10.4. The plate was incubated in the dark for 30 minutes, and then the reaction was stopped by adding 50 μL 3M NaOH. Absorbance was measured at 405nm wavelength and IgG concentration was calculated using an IgG calibration curve.

### Bacterial strains

*E. faecium* 11236/1, *S. aureus* MW2, *E. faecalis* T2, were obtained from Dr. von Hauner’s Children’s Hospital strain collection.

### Recombinant protein production

Recombinant AdcA and ZnuAdomains were produced as controls in OPA and OPIA assays, as previously reported (31). Briefly, the AdcA gene was amplified using the primers in Table S1 and digested with restriction enzymes BamHI/PstI (New England Biolabs). Amplicons were inserted downstream of the IPTG-inducible promoter into the pQE30 expression vector (QIA expressionist kit; Qiagen). The construct was heat-shock-transferred into the *E. coli* TOP10 (DE3) and *E*. *coli* BL21 (DE3). The ZnuA gene was amplified with primers in Table S1 and amplicons were digested and inserted downstream of the IPTG-inducible promoter into the pQE30 expression vector. The resulting construct was electroporated into *E*. *coli* M15pRep4. Recombinant ZnuA was purified in denaturing conditions using the Protino Ni-NTA Agarose (Macherey-Nagel) resin, following the manufactures instructions. Purified proteins were desalted by diafiltration using the Amicon Ultra-15 Centrifugal Filter Units of 3KDa (Merck-Millipore) and dialysed to a buffer composed of 150mM NaCl, 50mM Tris 7.8, and 2.5% (v/v) glycerol, for immunisation experiments.

### Opsonophagocytic Killing Assay (OPA) and Opsonophagocytic Inhibition Assay (OPIA)

For OPA and OPIA assays, we used *E. faecium* 11231/6, *S. aureus* MW2, and *E. faecalis* T2, which were grown in tryptic soy agar (TSA) and broth (TSB) at 37 °C. *S. aureus* MW2*, E. faecium,* and *E. faecalis* were cultivated in TSB, *S. aureus* with and enterococci without agitation. Full protocols of both assays were previously described elsewhere(42). Shortly, four components were prepared: (i) baby rabbit absorbed with the target bacterial strain as a source of complement, (ii) rabbit sera before and after immunization with Q11-EH and recombinant AdcA, (iii) polymorphonuclear neutrophils (PMNs) freshly prepared from human blood collected from healthy adult volunteers, and (iv) the bacterial strains grown to OD600 =C0.4 in tryptic soy broth (TSB).

For the OPA assay, the four components were mixed: 100 µL of PMNs (2.5×10^4^ µL^−1^); 100 µL of the appropriate serum concentration, 100 µL of complement (1∶15 dilutions for *E. faecium* and *E. faecalis*, and 1∶30 for *S. aureus)*, and 100 µL of an appropriate dilution of bacteria (1:200 *E. faecium, E. faecalis* and 1:75 *S. aureus*). All dilutions were made in RPMI supplemented with 15% FBS. The mix was incubated on a rotor rack at 37°C for 90 minutes, and after this time samples were diluted 100x in TSB and plated on TSA plates in quadruplicate. The percentage of killing was calculated by comparing the colony counts at 90 min of a control (bacteria in RPMI, no complement, cells, or antibodies) to the colony counts of a tube that contained all four components of the assay using the following formula:

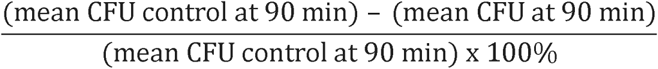

For OPIA assays, sera were diluted to a final concentration of 0.4mg/mL in RPMI + 15% FBS and pre-incubated overnight at 4°C with an equal volume of RPMI-F with either Q11-EH or AdcA, ZnuA(inhibitors) at final concentrations ranging from 3.56 to 0.14 µM.

Inhibition assays were performed at serum dilutions yielding 50–60% killing of the inoculum in the absence of inhibitors. Inhibition percentage was computed as the percentage of inhibition of opsonophagocytic killing with respect to controls without inhibitors. As controls, we used bacteria incubated in RPMI-F, in RPMI-F with complement at the corresponding dilution, in RPMI-F with PMN’s, and in RPMI with complement and PMN’s. Furthermore, to exclude minimal natural response in rabbits, sera collected before antigen administration were used as negative controls.

## DISCUSSION

Nosocomial infections (NI), acquired in healthcare units or after healthcare services, usually appear 48h after admission, or within 30 days after the patient’s discharge (43). The risk of NI is associated with the patient’s age, underlying diseases, length of hospitalization, immune system, or presence of invasive medical devices (44). Although employment of the mentioned devices may increase patient’s susceptibility to NIs, their use is often necessary for proper treatment and recovery. Multiple studies were undertaken to better understand and minimize the impact of AMR on healthcare (45, 46). The increasing prevalence of ESKAPE pathogens has emerged as a critical issue, especially in hospitals and long-term care facilities. AMR-NIs impose a substantial economic burden worldwide and significantly impact the quality of life and well-being of patients across all ages and genders (14, 15), with a higher impact in developing countries (4). Several candidate vaccine antigens have been proposed against *E. faecium* (23–27) and for *S. aureus* several clinical trials have been performed (29, 30) although none of these have shown promising results, and several explanations have been proposed to explain these failures (47). The development of a protective, effective vaccine against these pathogens, ideally cross-protective, remains an unmet need (28).

In our search for vaccine antigens with potential use against multiple pathogens, our approach involved the use of promiscuous smaller peptide sequences to identify an antigen capable of covering a diverse spectrum of pathogens (31). This “pan-vaccinomic” strategy would be a step toward a universal vaccine candidate, which could decrease the number of injections, facilitating vaccinations and increasing their accessibility. The antigen Q11-EH, which we present here, forms long fibrils exposing multiple copies of the EH-motif, which we previously have shown to act as a cross-protective antigen (31). Q11-EH was designed based on previous information that self-assembling peptides used as scaffolds for tissue engineering can be used as adjuvants (34). In physiological conditions, they can self-organize in long fibrils. These self-assembled peptide biomaterials have been shown previously to be well tolerated *in vivo* (37, 48) and it was also shown in mice that even upon administration of the self-assembling peptide Q11 with complete Freund’s adjuvant, no response toward fibrils was developed (34). Our data show that Q11-EH preserves the formation of nanometric fibrils, as observed for the Q11 peptide Q11 (34). With the Q11-EH supramolecular structure, we aimed to mimic repetitive molecular pattern presentation, which naturally occurs on the surface of pathogens such as bacteria or viruses, where antigenic proteins are expressed in multiple copies. Moreover, the large size of Q11-EH should facilitate the activation of a strong immune response (49). An additional advantage of our design was the flexibility of the antigenic sequences once exposed by the presenting fibre. Indeed, antigen flexibility is known to improve the immunological response, as it increases the variability of antigenic conformers during the recognition by B-cells and broadens the diversity of antibodies in the immune response (50). We have demonstrated that anti-Q11-EH antibodies successfully lead to bacterial opsonic killing in OPA assays with strains of *E. faecium*, *S. aureus,* and *E. faecalis* with greater efficacy compared to the full-length AdcA. The identification of an effective, albeit simple, self-assembling and multi-targeting antigen can be a step forward towards a universal vaccine antigen candidate to address AMR pathogens from the WHO concern list (51).

## AUTHOR CONTRIBUTIONS

R.B., F.S. and E.K. designed the peptide, developed the polymerization protocol and conducted DLS analyses. E.K., F.R., F.S., and R.B. prepared graphical content. D.M. and S.L.M synthesized Q11-EH and performed RP-HPLC and LC-MS analyses, while F.S. conducted ThT assays. F.R. selected rabbits for immunizations and, together with E.K., quantified IgG levels. E.K. and F.R. designed and executed OPA and OPIA experiments for *S. aureus* and *E. faecium*, with O.S. performing OPA experiments for *E. faecalis T2*. E.K. and F.R. performed statistical analyses, while E.K., F.R., R.B., and J.H. interpreted the data. The final version of the manuscript was prepared by E.K., and R.B. and edited by F.R., and J.H., with input from all authors.

## ACKNOWLEDGMENTS

Funding was provided by the project INF-ACT “One Health Basic and Translational Research Actions addressing Unmet Needs on Emerging Infectious Diseases PE00000007”, PNRR Mission 4, EU “NextGen-erationEU”- D.D. MUR Prot.n. 0001554 of 11/10/2022. E.K. and O.S. were funded by BactiVax - Anti-Bacterial Innovative Vaccines, Marie Skłodowska-Curie Actions, GA 860325. ). F.S. was supported by MUR through the PRIN2020 CANNOT-ESKAPE (2020XNFH9R): Targeting baCteriAl cell eNvelope of Nosocomial paThogens to ESKAPE resistance, 2021-2024.

