## Supplementary figures for "A self-assembling cross-protective antigen against multiple Gram-positive nosocomial pathogens"

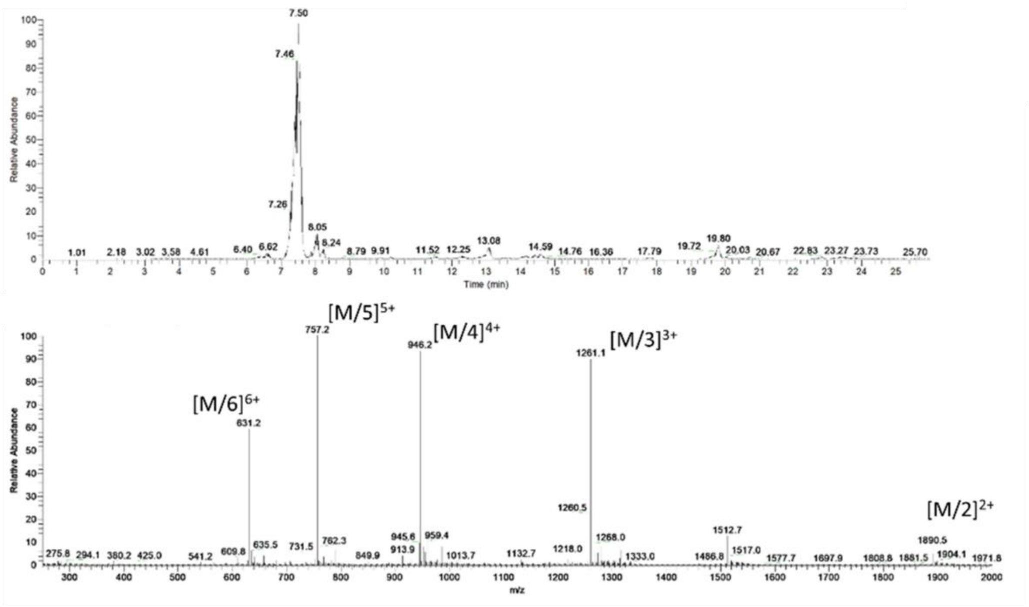


**Supplementary Figure 1.** Upper panel: LC-MS profile, Lower panel: MS spectrum of the main peak of crude non-acetylated Q11-EH peptide.

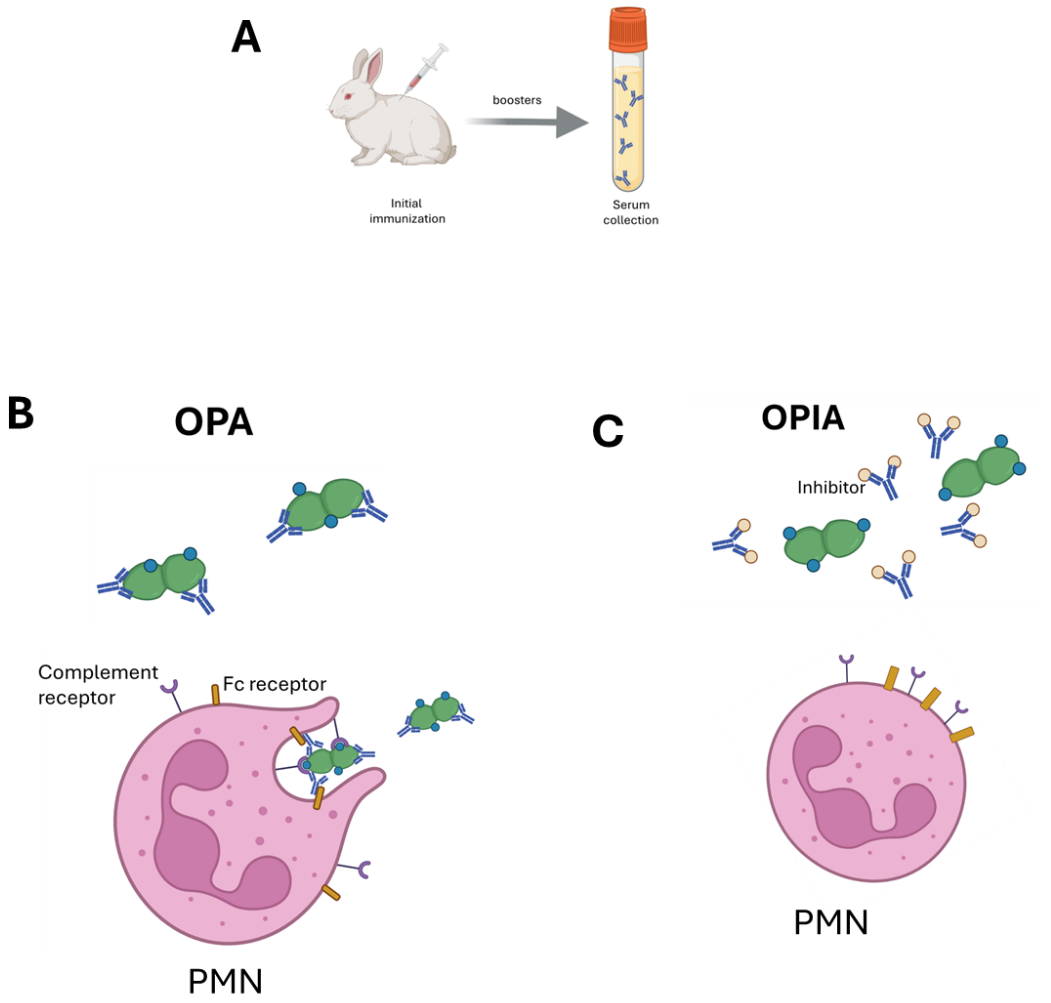


**Supplementary Figure 2.** A schematic representation of rabbit immunization (A) and mechanism of OPA (B) and OPIA (C). In panel B, antigen-specific antibodies and complement proteins opsonise bacteria and facilitate the uptake of the antibody-bacteria complex by phagocytes (PMN). In panel C, the inhibiting antigen binds antibodies, thus hampering opsonophagocytic killing.

**Supplementary Table 1.** Primers used for the recombinant production of AdcA and ZnuA domain

| | |
|---|---|
| AdcA | AdcA-5-BamHI aggcGGATCCTCGAATGATAAAGATGGAAAAT<br><br>AdcA-3-PstI aggcCTGCAGTTAATGAGCCATCATTTCTTGA |
| ZnuA | actacGGATCCAAATTAGAAATTGTAACAAC<br><br>gactagAAGCTTAAACCATAGTTGTTTTTCTAACGC |
